# Optimal coupling and task-specificity when learning rhythmic synchronization with a tool with varying levels of predictability and controllability

**DOI:** 10.64898/2026.04.02.716172

**Authors:** Dobromir G Dotov, Harjo de Poel, Claudine Lamoth

## Abstract

Sensorimotor learning and tool use involve synchronizing with external dynamics. Many everyday tools possess nonlinear hidden dynamics. Here we investigate how learning to synchronize with the complex dynamics of a tool depends on the degree of predictability and reciprocal coupling between user and tool. We introduce the concept of optimal coupling to measure adaptive user-tool coordination. Groups of participants practiced tracking an auditory stimulus in three conditions: 1) the tool was non-interactive and produced a periodic stimulus, 2) non-interactive and unstable stimulus, and 3) unstable but interactive stimulus which was coupled weakly to the participants movements and thus afforded control. Learning, retention, and transfer to visual modality were assessed using unpracticed test stimuli. Directional effective coupling was quantified using transfer entropy. Results showed that learning tended to be task-specific and there was no transfer to the visual modality. Interactive unstable practice exhibited some retention and generalization. We found a convergent reorganization of coupling during practice with the interactive unstable tool: stimulus-to-human coupling started high and decreased while human-to-stimulus coupling started low and increased. This suggests that embodiment of personalized rehabilitation technologies brings optimal reciprocal coupling in which sensorimotor-tool control is consistent with the minimal intervention principle postulated for within-body control.

---

Sensorimotor skills are fundamental for human action. We can move our bodies and manipulate objects in irregular and unpredictable environments. This often involves using elements of the environment as tools or interacting with machines which extend our functional reach and expand what we do. Studying the acquisition, loss, and re-learning of such skills is essential for improving human independence and quality of life. Here, we consider sensorimotor learning and complex tool acquisition as partially overlapping domains, defined as *sensory-motor-tool* learning. To address sensory-motor-tool learning in its full complexity, we need to consider that tools can have their own dynamics. Coupling of user and tool dynamics naturally creates a potential for nonlinear instabilities and unpredictable behavior. This can be a challenge for learning but may also support the development of more robust and flexible skills.

A popular approach to motor (re)learning consists of synchronizing with a repetitive target stimulus. This has shown promise in the cueing of gait in Parkinson’s disease and post-stroke (Awad et al., 2024; Roerdink et al., 2007, 2011; Thaut et al., 1996; Yoo & Kim, 2016), re-training of upper-limb mobility and dexterity in post-stroke (Schaefer, 2014; Van Vugt et al., 2014), and measuring sensory-motor synchronization abilities (Dalla Bella et al., 2012; Repp & Su, 2013), among others. Repeated tones or music with a fixed auditory rhythm support functional task performance in different clinical populations such as people with Parkinson’s disease or stroke survivors (Rochester et al., 2005; Spaulding et al., 2013), for example by providing continuous external timing signals to facilitate movement coordination and enhance gait (Dvorsky et al., 2011; Ploughman et al., 2018).

Synchronization tasks in which the stimulus is pre-recorded and fixed can be described as a one-directional coupling scenario in which the learner’s dynamics is coupled to the stimulus but not the other way around. In this context, effectiveness depends fully on the individual’s abilities for auditory-motor synchronization and beat perception (Dalla Bella et al., 2018; Zagala et al., 2024). In contrast, bi-directional coupling where the stimulus responds to the movements of the learner can personalize the stimulus and promote spontaneous synchronization across different skill levels. In these closed-loop tasks, high-frequency sensors and mobile computing allow the stimulus to continuously adjust its tempo and phase to match the participant’s tempo and phase (Dotov et al., 2019; Hove et al., 2012; Miyake, 2009; Moens et al., 2017; Repp & Keller, 2008; Szydlowski et al., 2019; Uchitomi et al., 2013). This approach may be particularly advantageous for rehabilitation because it more closely approximates the real-world interaction with the environment than a one directional coupling scenario.

Another way of increasing the relevance of rehabilitation tasks is to use target stimuli with less predictable dynamics. Synchronization with isochronously rhythmic predictable stimuli is convenient for starting to train sensorimotor and timing skills, but it does not capture the interaction with complex tools that possess their own dynamics. It remains unclear whether rhythmic synchronization generalizes to interactive contexts. Thus, fixed predictable stimuli may be suboptimal for rehabilitation, which typically requires not only accurate and precise motor timing but also skillful manipulation of tools and interaction with machines.

## Stabilization of unstable nonlinear dynamic systems as a generic paradigm for learning to interact with complex tools

Human-machine interaction often involves underactuated tools with internal degrees of freedom. This means that the user influences some degrees of freedom only indirectly through their coupling to other degrees of freedom. Manipulating underactuated tools, a fundamental aspect of many activities of daily living, can therefore elicit nonlinear phenomena, including unpredictability, instability, and structured variability. For example, carrying a cup of water without spilling requires active control to coordinate hand forces and motions so that the high-dimensional, nonlinear fluid dynamics are constrained to a predictable pattern (Sternad & Hasson, 2016). One way to achieve this is to avoid over-powering the object and instead exploit a stable periodic mode intrinsic to the object dynamics. This suggests that the Bernstein-inspired principle of minimal intervention, widely applied in theories of motor control and learning (Todorov & Jordan, 2002), may also apply to the manipulation of complex tools (Kant, 2016). This stresses the need to investigate the learning of nonlinear control and synchronization with complex rhythms. For example, participants can interact with richly dynamic stimuli in virtual space as if they are playing a computer game (Nayeem et al., 2023; Sternad, 2015).

Interaction with parametrically controllable complex tools has the potential to address another important target of rehabilitation strategies, namely, to generate movement pattern variety with personalized level of difficulty. Movement is inherently variable and therefore interventions should be designed to elicit *optimal* variability in tasks such as gait and postural control. (Kimijanová et al., 2024; Stergiou et al., 2006; Stergiou & Decker, 2011; van der Kooij et al., 2001). Adding noise to the target stimulus may encourage movement variability, but it may also obscure the underlying fixed predictable structure. A more versatile principle is to employ an active stimulus that generates structured irregularity due to dynamic instabilities in its internal degrees of freedom. A general model of this scenario is given by chaotic systems that are *deterministic*, in that they consist of a small set of nonlinear dynamical equations, yet *unpredictable* and *noise-like*, in that their trajectories do not exactly repeat.

Another remarkable feature of chaotic systems is that they can be controlled despite their apparent nonlinearity and unpredictability. For this reason, the mathematical framework of chaos control can be exploited to study how humans learn to interact and synchronize with complex tools. We used a chaotic system as the generative model for the stimulus and experimentally manipulated the coupling strength and unpredictability by setting parameters of the generative model (Dotov & Froese, 2018). This is a general experimental template that applies both to sensorimotor synchronization and complex object manipulation. This template can compare the effectiveness of different strategies for skill development by setting parametrically the unpredictability and reciprocal coupling, i.e. interactivity of the stimulus.

At the mechanistic level, chaos control consists of stabilizing a chaotic system by exploiting its deterministic structure. This is possible because the attractor set of a chaotic system is composed of unstable periodic orbits (UPO’s). A driving signal that is appropriately timed when the chaotic trajectory passes close to an UPO can stabilize the trajectory on the UPO, a phenomenon also known as chaos entrainment (Ott et al., 1990; Pikovsky et al., 2003; Rosenblum et al., 2003). This is comparable to balancing a multi-link or flexible inverted pendulum by driving it into a low-amplitude oscillation. For this to be possible, however, the driving signal needs to anticipate the unstable manifolds of the driven system, implying some form of learning. This likely requires first probing the system by applying perturbations in different directions and observing its responses (Boccaletti et al., 2002). In short, the same chaotic system can produce unstable oscillations when it is in the autonomous regime (*unstable noninteractive*) or become amenable to entrainment control when it is in the interactive regime (*unstable interactive*), provided that the driver knows how to push it. Furthermore, other parameters allow to reduce the unpredictability of the generative model to convert it to a *periodic noninteractive* stimulus.

## Measuring effective user-to-tool and tool-to-user coupling

The experimental template of chaos control emphasizes also that measuring the directional effective coupling from tool to user and from user to tool is necessary to understand task performance. In interactive tool scenarios, synchronization does not unambiguously indicate which part of the human-machine pair is contributing more to achieving overall performance as each member has some capacity to track the other member. In fact, quantifying the *effective* coupling in human-machine interaction is a difficult and often overlooked problem (Dotov et al., 2010; Segil et al., 2022). Measuring changes in effective coupling occurring with learning is needed to answer questions such as which member adapts as performance improves, what is the minimal parametric coupling that challenges performance for the sake of rehabilitation, and whether symmetric or asymmetric coupling better supports rehabilitation. Importantly, effective coupling can be distinguished from *structural* coupling that reflects the availability of sensor signals rather than the strength of functional interaction. Historically, effective coupling has been linked to the notion of information transfer between interacting entities.

The present study tested whether practice variability induced by interacting with an unstable nonlinear but controllable dynamic system can enhance the generalization of learning. We varied and extended previous work (Dotov & Froese, 2018) by adding retention and transfer testing. Different models generated the target stimulus in different practice conditions and pre, post, and retention tests. To probe transfer, one of the test tasks switched from the auditory to the visual domain. Participants’ basic task was comparable to playing a simplified hand-held monophonic musical instrument along with an artificial partner, the goal being for the two to play in perfect unison. To match the rhythm and pitch of their partner, participants had to control their hand movements and/or learn to control the partner through their hand movements. In transfer trials, the stimulus was a target object moving laterally on the computer screen, and the hand-held controller no longer modulated a sound but moved a tracking object on the screen.

## Methods

### Participants

Thirty young healthy adults volunteered. Exclusion criteria were hearing or motor control problems. Twenty-six participants (*N* = 26) completed the full study by attending the laboratory in two days. The study was approved by the Central Ethical Review Committee of the University Medical Center Groningen.

### Design and Procedure

Participants were assigned to one of three practice conditions. Practice consisted of 40 trials, each one minute long. After each practice trial, participants received intrinsically rewarding feedback in the form of a graph of their performance score history. Learning and transfer were assessed using four test tasks administered before practice (Pretest), immediately after practice (Posttest), and on a separate day (Retention).

Depending on the testing or practice condition, the target produced either periodic, Figure 1A or chaotic oscillations, Figure 1B-C. In the latter case, the oscillations had unpredictable changes in amplitude and period. Stimuli based on chaotic systems were labeled *unstable* because chaos entails at least one positive Lyapunov exponent (a positive divergence rate). This terminology distinguishes our paradigm from motor learning approaches that treat variability as additive noise. To study learning with unstable, hidden nonlinear dynamics, we implemented the *unstable interactive* task, in which the chaotic stimulus was weakly driven by the participant’s hand movements. This closed loop allowed participants to learn to stabilize the stimulus.

**Figure 1.**
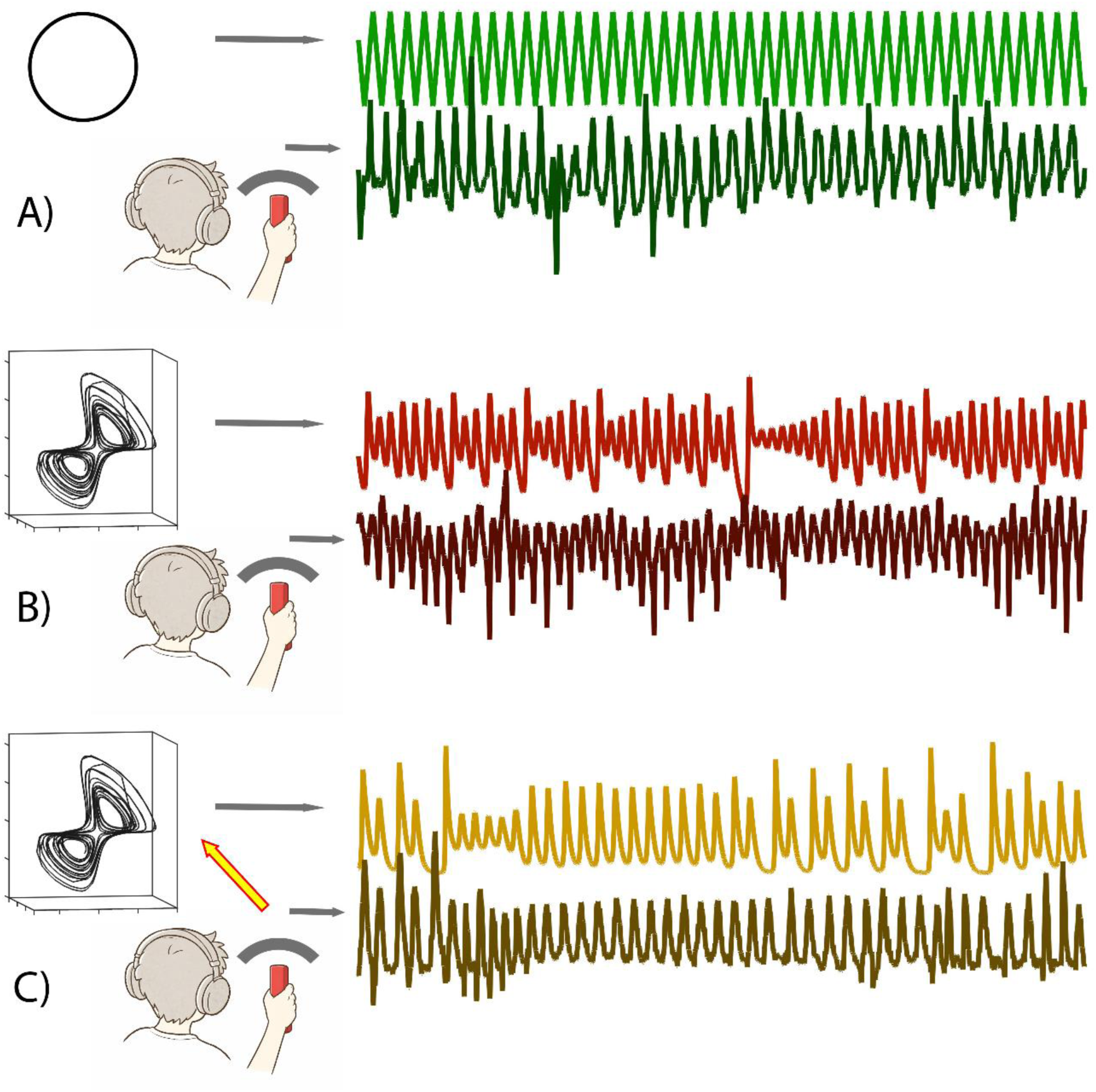
Schematic of the task space in each group and representative time-series from the stimulus and participant movement. A) a non-interactive (autonomous) phase oscillator generating a periodic stimulus, B) a non-interactive (autonomous) Chua attractor generating a chaotic stimulus, C) an interactive (non-autonomous) Chua attractor weakly coupled to the participant’s movements and generating a chaotic stimulus with the potential to be driven into a periodic orbit. In this example, it was held close to a one-cycle period temporarily and then a two-cycle period. A performance score accounted for both spatial (pitch-matching) and temporal (rhythm synchronization) precision by computing the average windowed cross-correlation between the two timeseries and dividing it by the root-mean-squared deviation of the participant from the stimulus.

#### Task

Across tasks, the overall goal was to match the pitch of the participant-generated continuous sound to the pitch of the target stimulus sound (see Figure 1). This was described figuratively as playing a musical instrument in unison with a partner who is the lead. In the transfer task, the sounds were replaced by visual target and tracker objects oscillating laterally on the computer screen.

#### Practice conditions

- *Periodic noninteractive.* The target was a fixed oscillation consisting of a non-interactive (autonomous) phase oscillator generating a periodic modulation of the stimulus pitch (Figure 1A).
- *Unstable noninteractive.* The target was irregular oscillatory modulation of the stimulus pitch. The oscillation was generated by mapping one of the three dimensions of a chaotic Chua oscillator to the stimulus pitch (Eq. 1).

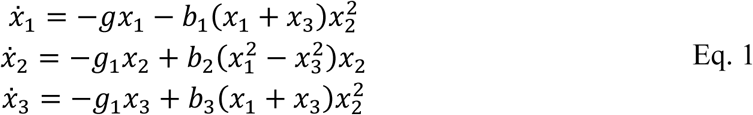 Here, *g*=-1, *b*_1_=9, *b*_2_=5, *b*_3_=1, and the driving force was given as parametric modulation *g*_1_=*g*_0_ – *u* with *g*_0_=1 (Chowdhury et al., 2001). As this was a noninteractive condition, the driving input *u*=εθ_x_ was zeroed and the generative system was in autonomous mode by setting the coupling parameter ε=.0 (Figure 1B).
- *Unstable interactive.* The target stimulus was the same way as in the unstable noninteractive condition (Eq. 1), but the generative system was interactive (non-autonomous mode) by setting ε=.7. The input signal θ*_x_* was the vertical tilt of the participant’s hand-held controller. This weak coupling with the participant’s movement made it possible to stabilize the stimulus on a periodic orbit (Figure 1C).

In sum, we manipulated the controllability and predictability of the target generator. Conditions differed in their affordances for synchronization, variability-generating instability, and reciprocal control, which together defined a multidimensional parameter space. Each condition occupied a distinct region in this space. The *periodic noninteractive* stimulus afforded high synchronization but low controllability and required minimal reactive adjustment. The *unstable noninteractive* stimulus was difficult to synchronize with, contained challenging instabilities, elicited corrective responses, and did not afford control. The *unstable interactive* stimulus was similarly difficult to synchronize with, but it responded to participant movement, affording control and exploration of its intrinsic dynamics. To compare learning, retention, and transfer of the rhythmic sensorimotor tool skill between the three practice conditions, we used the following variations of the task in pre, post, and retention tests.

#### Test and transfer tasks

- Test 1: *Periodic noninteractive test* with stimulus with different amplitude and frequency differing from those used in practice
- Test 2: *Periodic noninteractive continuation test* in which the stimulus disappeared after a few seconds and participants maintained its period and amplitude.
- Test 3: *Unstable noninteractive test* using one dimension of an unpracticed chaotic system (Lorenz attractor) running without participant input (autonomous mode).
- Test 4: *Transfer unstable noninteractive* test using the same target stimulus as Test 3, displayed as a white visual target moving left-right on a black screen.

### Apparatus

The experiment was performed using a desktop computer and screen, headphones, and a hand-held wireless game controller, while sitting at a desk in a quiet lab space. The motion sensing device was a Nintendo Wii controller with embedded accelerometer and Bluetooth link to the computer. A custom Python program processed in parallel the sensor signal, the target stimulus, and sound and visual synthesis. The stimuli were generated in real-time using ordinary differential equations integrated numerically in steps each time the sensor sample was updated. The Wii controller was used as a tiltmeter by mapping its inclination in the range from -90 (left) to +90 (right) degrees (-1 to +1 g) to auditory pitch of a pure tone in the chromatic scale. The sensor was sampled at a varying rate between 90 and 100 Hz.

The sounds of the participant’s controller and the stimulus were synthesized in real time at 48 kHz. There were two synthesizer instances running in parallel, corresponding to the left and right channels. Each synthesizer produced a sustained pure tone with two harmonics while its pitch was modulated either by the participant or the stimulus, respectively. Participants heard the stimulus on the left side and themselves on the right side, which was the side holding the controller, unless the participant requested left-handed mode. We used the same sonification engine as in a previous study because it was designed with the beneficial properties of being fast, efficient, and capable of automatically scaling amplitude and frequency to roughly preserve equal loudness within the lower speech range of hearing (Dotov & Froese, 2018).

### Measures

#### Task performance score

To account for two important aspects of task performance, rhythm matching and pitch matching, we computed a composite score, synchronization divided by pitch error. Specifically, the average absolute windowed cross-correlation between stimulus and participant trajectories for each trial was divided by the root-mean-squared deviation between the two trajectories.

#### Effective Coupling

Directional coupling between participant and stimulus was assessed using transfer entropy (TE), a directional conditional mutual information between source system *X* and target system *Y* (Schreiber, 2000). *TE* is a directional conditional mutual information between source system *X* and target system *Y*, represented by time-series *x* and *y* which are delay-embedded in *d*-dimensional vectors 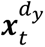 and 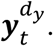.

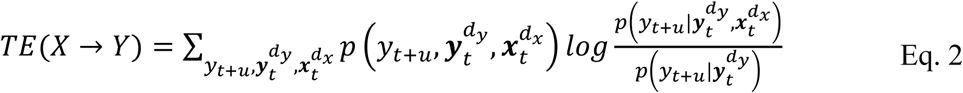

In practice, Shannon entropy is used.

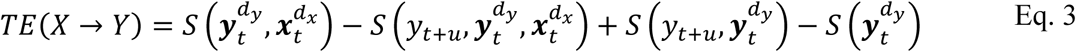

In the original definition, *u* is equal to one time step but it can be optimized for the coupling between the given pair of systems (Wibral et al., 2013). *TE* is an alternative to Granger causality.

For each trial, we estimated transfer entropy from the stimulus to the participant, *TE*_Stimulus→Human_, and from the participant to the stimulus, *TE*_Human→Stimulus_. The former was interpreted as the strength of the participant’s reactive responses to the stimulus oscillations. The latter was interpreted as the participant being able to control the stimulus. *TE* was estimated using the TRENTOOL toolbox (Lindner et al., 2011) with fixed embedding, *m*=3 for chaotic and *m*=2 for periodic stimulus trials. To control for finite-sample effects and superficial similarities between time series, we used a surrogate approach that preserve statistical properties of the data but destroy any coupling linkage. For each task, we constructed a surrogate distribution by pooling trials across participants and reassigning one member of each paired time series to a different trial, repeated three times. *TE* values were rescaled as *z*-scores relative to this surrogate distribution. A rescaled *TE* score of 2 indicates two standard deviations above the surrogate mean.

#### Instability

Instability was quantified via the maximum Lyapunov exponent λ, reflecting attraction (negative exponents) and repulsion (positive exponents) along the attractor’s manifolds. We computed the maximum Lyapunov exponent of hand and stimulus oscillations using the Rosenstein algorithm (Rosenstein et al., 1993). In human movement, this method has been applied to gait (Bruijn et al., 2013; Ijmker & Lamoth, 2012; Miller et al., 2006). Also called log-divergence, λ quantifies divergence of adjacent trajectories in reconstructed phase space. We obtained both long-term divergence at the trial timescale and short-term divergence within a single movement cycle. Recorded hand and stimulus trajectories were low-pass filtered (6 Hz). Hand movement was delay-embedded into a three-dimensional phase space with delay equal to a quarter of the average period. For consistency, the same delay-embedding method was applied to the periodic stimulus. For stimuli in unstable trials, phase space reconstruction was skipped because the full system was already available.

### Analysis

With *N* = 26 valid participants across three groups, analyses were treated as exploratory. We tested for improvement relative to the pre-test in each group and each test task separately without correcting for multiple comparisons. For each test task, multiple trials were averaged, and for each measure, percentage improvement from pretest to posttest, and from pretest to retention was calculated. One-sample *t*-tests evaluated the null hypothesis of zero improvement. Paired-sample *t*-tests evaluated the null hypothesis of no change from post-test to retention.

Practice data were analyzed separately from test data using linear mixed-effects models to capture improvement across practice trials, with the periodic noninteractive practice group as the reference condition. Performance was analyzed using a linear mixed-effects model with trial number as a time-varying predictor, practice group as a between-subjects categorical predictor, and a trial-by-group interaction term. Dependent variables were performance score, *TE* in the human-to-stimulus and stimulus-to-human directions, and short and long-term Lyapunov exponents of the stimulus and the participant. *p*-values <0.05 were considered significant.

## Results

### Performance and dynamic parameters in practice trials

As Figure 2 shows, each practice condition was characterized by its distinct dynamic signature of dynamic instability and affordances for synchronization and control.

**Figure 2.**
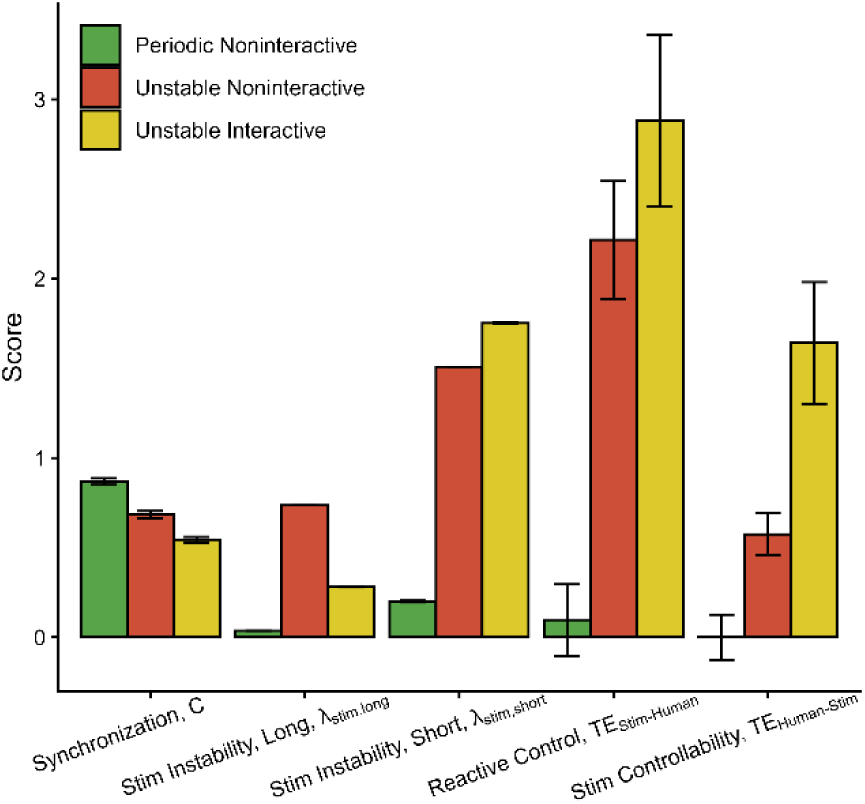
The unique dynamic signatures of the three practice tasks can be illustrated by parameterizing their dynamic instability and affordances for synchronization and control. Bars are *Mean*±*SE* measured in the practice trials in this study. Synchronization was measured with the windowed cross-correlation. The instability of the stimulus at the trial scale was measured with the long-term maximum Lyapunov exponent. The instability of the stimulus at the cycle scale was measured with the short-term maximum Lyapunov exponent. For reactive control exercised by the participant, we measured effective coupling using the stimulus-to-participant transfer entropy. For controllability of the stimulus, we measured effective coupling in the opposite direction using the participant-to-stimulus transfer entropy.

#### Taks Performance Score

Performance improved across the 40 practice trials, as indicated by a significant effect of trial number (β = .0036, *t* = 4.74, *p* < .001). Relative to the periodic noninteractive condition, overall performance was lower in both the unstable noninteractive group (β = −.287, *t* = −3.12, *p* < .001) and the unstable interactive group (β = −.296, *t* = −3.38, *p* < .01). The trial-by-group interaction was not significant (all *p*’s n.s.).

#### Effective coupling from stimulus to participant

Stimulus-to-human TE did not change across practice trials in the periodic noninteractive group (*p* = .21). In the unstable noninteractive group, stimulus-to-human TE was significantly higher than in the periodic group (β = 1.94, *t* = 3.50, *p* < .01) and remained stable across trials (*p* = .28). In the unstable interactive group, stimulus-to-human TE was also higher than in the periodic group (β = 3.48, *t* = 6.62, *p* < .001) but declined significantly over the course of practice (β = −3.40, *t* = −4.45, *p* < .01), indicating that participants became progressively less reactive to the stimulus as practice proceeded.

#### Effective coupling from participant to stimulus

Human-to-stimulus TE did not change across practice trials in the periodic noninteractive group (*p* = .34), nor did it differ from the periodic group in the unstable noninteractive group (*p* = .46). A marginal increase across trials was observed in the unstable noninteractive group (β = .015, *t* = 2.05, *p* = .04); however, this result should be interpreted with caution, because by design of the experiment, participants in this condition could not physically affect the stimulus, thus human-to-stimulus TE was expected to remain low throughout practice. Consistent with this, TE values in the unstable noninteractive group remained consistently below two standard deviations above the surrogate mean across all trials (Figure 3B).

**Figure 3.**
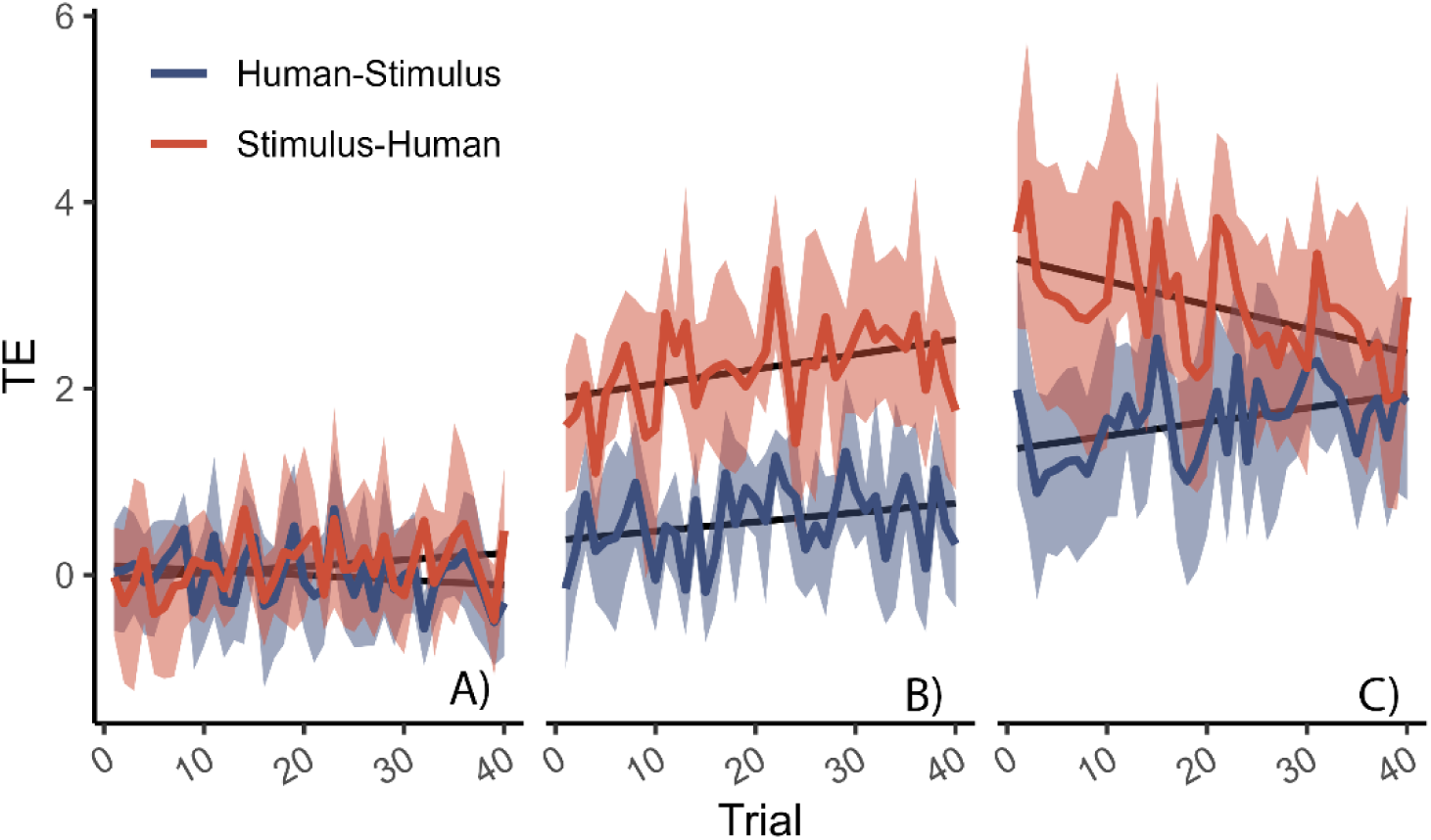
Change along practice trials (Mean±95%CI across participants) of the directional effective coupling. Coupling was calculated with the transfer entropy rescaled as the *z*-score relative to a surrogate distribution. A) Periodic non-interactive stimulus group. B) An unstable non-interactive stimulus group. C) Unstable interactive group.

In the unstable interactive group, human-to-stimulus TE was significantly higher than in the periodic group (β = 1.25, *t* = 3.57, *p* = .01) and increased across practice trials (β = .019, *t* = 2.77, *p* < .01), providing further evidence that participants in this group were learning to stabilize the stimulus. Notably, over the course of practice, stimulus-to-human and human-to-stimulus TE converged in the unstable interactive condition (Figure 3C).

#### Instability of the stimulus

As intended by design, the short-scale (subcycle) Lyapunov exponent of the stimulus was significantly higher in both the unstable noninteractive condition [β=.307, *t*=186.931, *p*<.001] and the unstable interactive condition [β=1.563, *t*=235.483, *p*<.001] compared to the periodic condition (see Figure 2). Stimulus instability did not change across trials in the unstable noninteractive condition [*p*=0.61]. In contrast, there was a significant trial by unstable interactive group interaction [β=-.0005, *t*=-2.542, *p*<.05], indicating that participants in the interactive condition progressively reduced stimulus instability over practice—confirming that they were learning to control the stimulus.

The same pattern was observed for the long-scale maximum Lyapunov exponent of the stimulus. Stimulus instability was higher by design in both unstable conditions [β=.700, *t*=324.584, *p*<.001; β=.249, *t*=121.471, *p*<.001] (see Figure 2), with no overall change across trials [*p*=0.91]. A significant reduction across trials was again observed in the interactive condition [β=-.0002, *t*=-2.790, *p*<.01]. For illustration, Figure 1 shows a representative trial with moderate degree of control where the participant managed to temporarily entrain the stimulus on periodic orbits but these were difficult to maintain.

#### Instability of hand movements

With respect to participants’ hand oscillations, the short-scale Lyapunov exponent was significantly higher in the unstable noninteractive group than in the periodic group [β=.362, *t*=11.22, *p*<.001], whereas the unstable interactive group did not different from the periodic group [β=.035, *t*=1.14, *p*=.26]. Neither group showed a significant change across trials [all interaction *p*’s n.s.]. The long-scale Lyapunov exponent of hand movements exhibited a different pattern (see Figure 4). The periodic group showed no change across trials [*p*=.71]. The unstable noninteractive group did not different from the periodic group at the beginning of practice [*p*=.10] but instability increased significantly over trials [β=.0001, *t*=2.676, *p*<.01] (see Figure 5), indicating that participants were learning to produce instability. Similarly, the unstable interactive group started with al lower long-term Lyapunov exponent, than the periodic group at [β=-.037, *t*=-2.87, *p*<.01] but also increased instability with trial, reflecting the same learning-to-produce-instability effect [β=.0006, *t*=2.978, *p*<.01], (see Figure 4).

**Figure 4.**
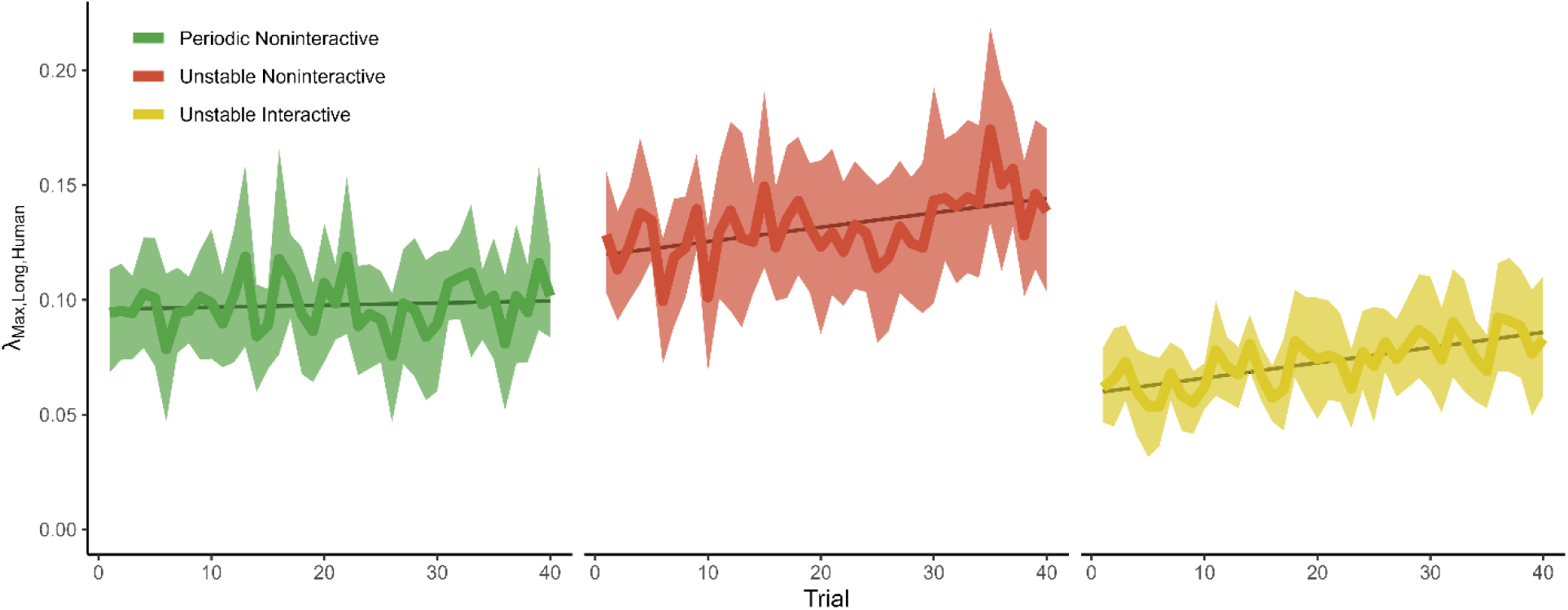
Instability of hand movements during practice trials (Mean±95%CI across participants). Instability on the time scale of multiple cycles was measured with the long maximum Lyapunov exponent. Groups are color-coded.

**Figure 5.**
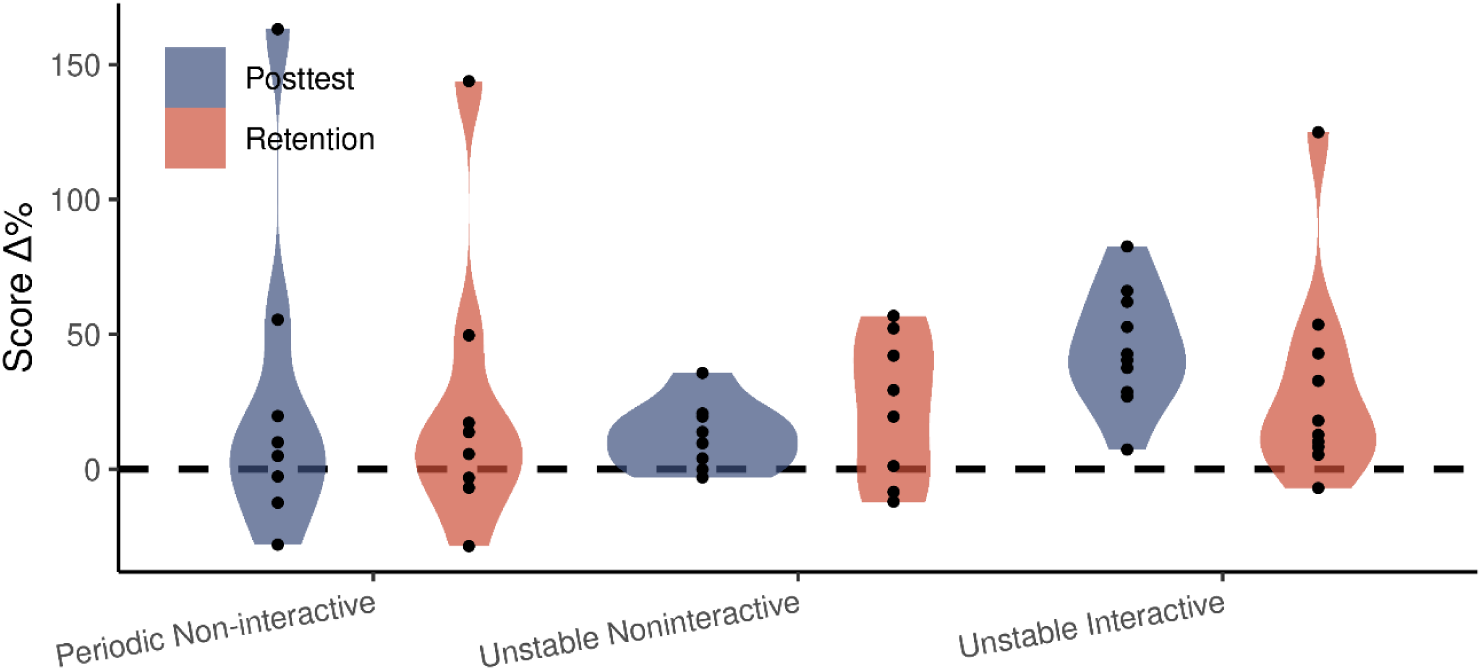
Testing for generalization of practice across practice groups. Test 3 was an unstable noninteractive test with an unpracticed stimulus generator, the Lorenz system.

### Performance improvement in post-test and retention

Periodic noninteractive test with stimulus with different amplitude and frequency (Test 1) did not show a significant improvement from pre-test to post-test and retention in any of the practice groups.

For the periodic synchronization continuation task (Test 2), the periodic noninteractive practice group improved significantly from pre to post-test [*t*(7)=3.86, *p*<.01] and this was also significant at retention [*t*(7)=5.14, *p*<.01]. There was no difference between post-test and retention [*t*(7)=-2.14, *p*=.07]. The unstable noninteractive group did not show a significant pre-test to post-test change [*t*(7)=1.43, *p*=.20] but, unexpectedly, showed improvement at retention [*t*(7)=2.95, *p*<.05]. There was no difference between post-test and retention [*t*(7)=-.73, *p*=.48]. The unstable interactive group improved from pre-test to post-test [*t*(9)=2.59, *p*<.05] but the pre-test to retention change was not significant [*t*(9)=2.02, *p*=.08]. There was no difference between post-test and retention [*t*(9)=-.35, *p*=.74].

For the unstable noninteractive task (Test 3) the periodic noninteractive practice group did not improve from pre- to post- [*t*(7)=1.23, *p*<.26] or from pre-test to retention [*t*(7)=1.27, *p*<.25] (see Figure 5). There was no difference between post-test and retention [*t*(7)=.25, *p*=.81]. In contrast, the unstable noninteractive practice group improved at post-test [*t*(7)=2.77, *p*<.03] but not at retention [*t*(7)=2.36, *p*=.05]. There was no difference between post-test and retention [*t*(7)=-1.37, *p*=.21]. The unstable interactive practice group improved, both at post-test [*t*(7)=6.46, *p*<.001] and retention [*t*(9)=2.51, *p*<.05]. There was no difference between post-test and retention [*t*(9)=.96, *p*=.36].

In the unstable noninteractive transfer task in a different sensory modality (Test 4), a visual stimulus was presented instead of an auditory stimulus. None of the practice groups showed significant improvement from pretest to posttest or retention [all *p*’s n.s.].

## Discussion

To describe the interactive task qualitatively, perfect stabilization was very hard to achieve because it required anticipation of the stimulus trajectory and very precise matching of its amplitude at the given moment. Because interaction with the stimulus was executed via an inertial sensing system, this was a complex task challenging rhythmic, spatial, and force control capacities. In practice, participants rarely succeeded in balancing the stimulus throughout the trial, just like balancing a flexible inverted pendulum is nearly impossible without extended training.

The present study investigated rhythmic sensorimotor-tool skill acquisition using a paradigm in which participants engaged in experimentally gated, real-time interaction with a dynamical system exhibiting nonlinear instabilities. The sensorimotor environment was built using sonificiation, the real-time mapping of movement to sound. Sonification can support motor rehabilitation because it capitalizes on the speed of low-level auditory processing, and can provide salient, low-distraction augmented feedback that supports timing and coordination in rehabilitation (Hermann et al., 2011; Scholz, 2015).

A key strength of this paradigm is its capacity for smooth hyper-parametric modulation of controllability and instability, enabling systematic comparison of regular (predictable) and irregular (unpredictable) rhythmic stimuli as well as interactive and non-interactive conditions, while holding other dynamic properties—including amplitude, frequency, and signal smoothness—constant across conditions. This design ensured that observed differences could be attributed to the manipulated variables rather than to incidental variation in low-level stimulus properties. The primary aim was to examine how bidirectional coupling between participant and stimulus shaped the *generalization* and *retention* of acquired motor skills, and how the nature of this interaction evolved across practice.

### Generalization and retention

To assess the extent to which learning transferred beyond the practiced stimuli, generalization and retention were evaluated using post-test stimuli that differed from those encountered during practice, including a cross-modal transfer task employing visual feedback. We found some, but modest, evidence for generalization as post-test and retention effects tended to be confined to the types of stimuli that the groups practiced with. Specifically, the periodic practice group improved in the periodic continuation post-test and retention. The unstable noninteractive group exhibited post-test improvement with a novel unstable stimulus but failed to retain this benefit at retention, suggesting that learning was present but did not stay at follow up. In contrast, the unstable interactive group, showed both post-test improvement and significant retention with a novel unstable stimulus, indicating more durable and generalizable learning. Both unstable groups showed some degree of improvement on the periodic continuation task, pointing to a degree of transfer from unstable to periodic stimuli. Neither group demonstrated improvement on the visual modality transfer task, suggesting that the acquired skills were tied to the trained sensory modality and did not generalize across modalities.

### Practice-related changes in movement instability

In practice trials, all three groups improved progressively over the course of practice trial, consistent with standard skill acquisition. Of particular theoretical interest were the group-specific changes in *instability* and directional *effective coupling*.

Participants in the unstable noninteractive condition increased their long-scale Lyapunov exponent across trials, reflecting an adaptive increase in the complexity of their motor output in response to an irregular, non-controllable stimulus. A qualitatively similar pattern emerged in the unstable interactive condition: participants likewise increased their long-scale movement instability over practice. Crucially, however, participants in this condition simultaneously learned to *reduce* the instability of the stimulus itself, an effect unique to the interactive group.

The most informative and least readily anticipated finding concerned the pattern of *effective coupling* as indexed by transfer entropy (*TE*). Stimulus-to-participant coupling was high in the two unstable conditions; however, it increased across practice for the noninteractive stimulus and decreased across practice for the interactive stimulus. Conversely, user-to-stimulus coupling increased in the unstable interactive condition. In fact, effective coupling in the two directions appeared to converge, see Figure 3C, and it converged towards a normalized *TE* value of about +2 *SD* which could be used as a pragmatic reference for meaningful or significant coupling.

### Optimal coupling with a tool

Effective coupling, also referred to as effective connectivity, is an important topic in the neurosciences and robotics, where questions about directional information flow between dynamical nodes are readily motivated (Friston, 2011; Lungarella & Sporns, 2006). Movement science deals with sensorimotor networks, yet it does not usually consider active handheld tools as integrated parts of sensorimotor control. The *tool embodiment hypothesis*, that functional tools are dynamically equivalent to parts of the body, is usually treated separately as a cognitive or a phenomenological question (Dotov et al., 2010; Froese, 2014; Segil et al., 2022).

Consequently, prior work offers limited basis for specifying a priori predictions about how effective coupling should evolve with practice with interactive stimulus in this study. Here, we demonstrate that directional coupling measures can provide an informative index of tool-mediated interaction. We use these results to suggest a conceptual framework to understand - changes in - effective coupling over practice.

What changes of effective connectivity are associated with the embodiment of a tool? One may be tempted to predict that practice in the unstable interactive condition leads to increased *TE* in both directions, consistent with higher information flow and better control. However work on dyadic human interaction, has shown that better synchronization can coincide with reduced effective coupling between two humans (Wood et al., 2022). Consistent with this possibility, we observed a pronounced initial asymmetry in directional coupling, with low human-to-stimulus *TE* and high stimulus-to-human *TE*, followed by convergence toward comparable values with practice (Figure 3C). Although this pattern may appear counterintuitive, yet it is consistent with the principle of minimal intervention (Todorov & Jordan, 2002), as control becomes more skilled, corrections are applied selectively, and unnecessary coupling is reduced. What is remarkable is that the principle intended for within-body sensorimotor control is found to apply just as well to tool control (Kant, 2016). Notably, the convergent pattern was driven in part by an increase in human-to-stimulus effective coupling with practice across trials. Because the stimulus parameters were not adaptively updated from one trial to the next, this change is unlikely to reflect altered structural coupling and instead suggests a practice-related change in how participants exploited the available interaction dynamics. Given that *TE* can be indicative of leader-follower relations (Wirkuttis et al., 2023), the convergence of *TE* between user and tool suggests that with practice the leader-follower division of labor was replaced by an optimally reciprocal interaction. To describe this mode of moderate but significant and reciprocal coupling in which the user and the tool make a fluently functioning system, we propose the notion of *optimal coupling with a tool*, loosely inspired by the phenomenological notion of *optimal grip* (Bruineberg & Rietveld, 2014).

Future work can exploit our formally grounded experimental task to advance the theoretical modeling of complex tool use. In the present study, we assumed that unstable periodic orbit (UPO) stabilization can serve as a paradigmatic model of learning to control a tool with internal degrees of freedom. Building on existing work on chaos control, a promising next step is to model how a simulated participant could be controlling the stimulus and compare to our empirical observations. Work in applied dynamic systems has discovered both model-based and model-free strategies for UPO control (Boccaletti et al., 2000). This is exciting because theoretical distinctions such as model-based versus model-free control and feedback versus feedforward control (Wichmann & DeLong, 2013) can be studied by testing how different existing models of chaos control explain human-stimulus interaction.

### Task specificity and Implications for rehabilitation

Our findings underscore the predominance of task-specific learning in rhythmic sensorimotor-tool skill acquisition. We adopted a dynamical systems framework to design stimulus generators with predictability and controllability that could be modified to instantiate different experimental conditions (Dotov et al., 2019; Dotov & Froese, 2018; Muratori et al., 2013). Despite theoretical and empirical prior work that variable and less predictable practice can support more persistent and generalizable learning (Merzenich, 2013). performance gains in the present paradigm were largely constrained to test stimuli that preserved the dynamical structure encountered during practice. Overall, the results showed that learning was largely task-specific: periodic practice improved performance on periodic test stimuli, and unstable practice improved performance on unstable test stimuli. We also observed some retention for the interactive practice stimulus, but no transfer to the visual transfer task.

From a rehabilitation perspective, the key issue is therefore not whether learning generalizes in the abstract, but which *type* of learning is required for a given clinical goal and which practice manipulations preferentially support it. Our principled approach could derive training strategies that may suit different learning requirements. When sensorimotor and tool skill learning depends on robust error-based calibration within a specific task and sensory modality, practice should prioritize that task and modality, and promoting errors through variability can be desirable (Schmidt & Bjork, 1992). In this study, practice with an unstable stimulus promoted the generation of increased instability by the participants. Given that induced instability in rhythmic dynamics is a mechanism for generating variability, our approach is promising for training beneficial variability when this is needed (Magill & Anderson, 2021).

There is no universally accepted taxonomy of motor learning types. However, several broad distinctions are widely recognized. For example, learning is often predominantly implicit (Wolpert et al., 2013), but it can also begin in explicit mode and gradually transition into implicit with repetition, such as when learning to drive a car or a bicycle (Magill & Anderson, 2021). In this study, participants were not instructed how to control the tool in interactive condition. At debriefing, they could not explain exactly what they were doing when they were succeeding, although this does not exclude the use of transient explicit strategies. The task also had elements of reinforcement learning because it provided an intrinsically rewarding trajectory of score improvement as feedback after each practice trial (Wichmann & DeLong, 2013).

In summary, our results suggest that sensorimotor skill with complex tools is shaped not only by the stability of the practiced dynamics, but also by the structure of interaction itself. We provided a framework which allows learning demands to be flexibly tuned through coupling and instability parameters. Beyond basic science, this flexibility could be exploited to design personalized applications and adapt practice difficulty over time. Thus, these parameters offer a principled way to individualize training and progression in neurorehabilitation, where the goal is often to challenge the system without inducing maladaptive compensation (Valero-Cuevas et al., 2024). We therefore view controllable instability and directional coupling metrics as complementary levers for designing interventions that are both theoretically grounded and clinically adaptable.

## Acknowledgments

We would like to thank Marije Harmsen and Liset van der Hulst for their indispensable help with this study.

## Notes

### Competing Interest Statement

The authors have declared no competing interest.

